# Levodopa induced dyskinesia severity is linked to distinct microstructural alterations in nigrostriatal and brainwide neural circuits

**DOI:** 10.64898/2026.10.30.763167

**Authors:** Mikhail Moshchin, Susan Osting, Emily Yakel, Roger J Schultz, Kevin P. Cheng, Jade E. Davis, Cuong P. Luu, James K. Trevathan, Wendell B. Lake, Samuel A. Hurley, Erin K. Purcell, Christopher Bishop, Kip A. Ludwig, Aaron J. Suminski

## Abstract

Dopaminergic replacement therapy via L-DOPA is the major medical therapy for Parkinson’s Disease. Long-term L-DOPA administration is known to cause dyskinetic movements, L-DOPA Induced Dyskinesia (LID). It remains unclear if the LID related, maladaptive changes in brain structure/function are localized to neural circuits within the nigrostriatal DA system or have brainwide impact. We employed multi-shell diffusion magnetic resonance imaging and correlational tractography to investigate structural brain networks associated with daily administration of L-DOPA and the development of LID in the unilateral 6-hydroxydopamine rat model. 53% of rats receiving L-DOPA therapy developed severe LID. Tractography found brainwide statistical maps positively and negatively associated with LID severity highlighted by bilateral neural circuits contained in the lesioned nigrostriatal system as well as others including the cerebellum and prefrontal cortex. These results suggest that systemic dysfuntion in brainwide neural circuits forms the mechanistic basis the maladaptive changes underlying LID.

## Introduction

Parkinson’s disease (PD) is one of the most prevalent neurodegenerative disorders, progressively impairing quality of life. Triggered by profound striatal dopamine deficiency, recent evidence points to PD being a network disorder characterized by structural and functional changes throughout broad action, motivation and arousal networks^1–4^. Accordingly, those with PD suffer from a debilitating array of motor (bradykinesia, rigidity, resting tremor, and postural instability) and non-motor symptoms such as sleep disturbances, depression, anxiety, pain, and fatigue^5^. Unfortunately, PD remains without a disease modifying therapy. Instead, symptomatic management via medical therapy or deep brain stimulation is available, with the current gold standard treatment being dopamine replacement therapy (DRT) with L-dihydroxyphenylalanine (L-DOPA). While highly beneficial in reducing motor deficits, long-term L-DOPA administration often leads to significant side effects like motor/non-motor fluctuations^6^ and severe dyskinesia^7^ (i.e. L-DOPA-induced dyskinesia (LID)). LID pathology involves widespread, maladaptive changes across multiple neurotransmitter systems and altered neural circuitry^8–13^.

This multifaceted etiology necessitates a paradigm shift in evaluating the mechanistic underpinnings of LID and approaches to mitigate/unwind the appearance of peak dose dyskinesia. Rather than strict focus on isolated targets like spiny projection neurons (SPNs) or single neurotransmitter systems^13,14^, a growing consensus advocates for a comprehensive neurocircuit-based perspective to unravel the mechanisms driving PD and LID. Work in both PD patients^1,15^ and animal models^16–18^, including our own^19,20^ demonstrate that PD is a whole brain disorder, and LID similarly involves widespread brain pathology/changes^21–28^. In particular, circuits involving the cerebellum and prefrontal cortex are modulated by PD, L-DOPA therapy^29^ and LID demonstrating their role in compensating for altered function of cortico-striatal-thalamo cortical networks. While these changes likely impact motor control circuits, the impact on nonmotor circuits cannot be underestimated given the known connectivity between the lateral cerebellum and prefrontal cortex^30^ and the symptom specific alterations in cerebellar prefrontal networks in early stage PD^20^.

To address this neurocircuit perspective on LID, we identified structural brain networks correlated with dyskinesia severity in a unilateral 6-OHDA rat model during chronic L-DOPA treatment, using a dose of L-DOPA known to reverse motor deficits, but also induce LID (6 mg/kg)^11,31^. This study aimed to characterize the structural changes induced by chronic DRT with L-DOPA, specifically contrasting those observed in the absence of abnormal involuntary movements (AIMs) with the neural networks underlying AIMs development. Using ex vivo, multi-shell diffusion magnetic resonance imaging (dMRI) and correlational tractography^32^, we assessed microstructural changes throughout the brain resulting from unilateral injection of 6-hydroxydopamine (6-OHDA) in the medial forebrain bundle (MFB) with and without chronic L-DOPA treatment. Consistent with reports on the development of dyskinesia following DRT in rodents^33^ and patients with PD^34^, our study revealed that 53% (8 of 15) of PD rats treated chronically with L-DOPA developed severe dyskinetic movements. Correlational tractography identified distinct, bilateral brain statistical maps both positively and negatively associated with both LID severity and disease state (False Discovery Rate [FDR] < 0.05). LID preferentially modulated nigro-striatal and cortico-striatial networks commonly associated with PD and LID but also those including the cerebellum and prefrontal cortex. Fiber tracts connecting the striatum to the substantia nigra pars compacta and pars reticulara were anatomically distinct and correlated with disease state and LID severity, respectively. These results highlight the emerging concept of broad network dysfunction driving symptomatic progression in PD and the maladaptive changes linked to the emergence of LID and provide a basis for mechanistic examinations of the network alterations in Parkinson’s progression and treatment response.

## Results

### PD Model: Behavioral and Histological Confirmation

To evaluate structural changes associated with long-term LID, we first established a unilateral PD model. On Day 0 (Fig 1A), we depleted dopamine by injecting 6-OHDA into the left MFB of 48 rats; 6 rats were injected with saline alone (Vehicle cohort). Fourteen (14) days post-surgery, we assessed each rat’s motor symptom development with the cylinder and apomorphine (APO) tests (Fig. 1A). During the apomorphine test, 6-OHDA injected rats that did not rotate contralesionally more than 7 rotations per minute (n = 26) were excluded^35^, leaving 22 rats for the treatment phase (PD cohort). Two rats in the Vehicle cohort were excluded from the study due to surgical complication and inadequate brain coverage during dMRI.

**Fig. 1.**
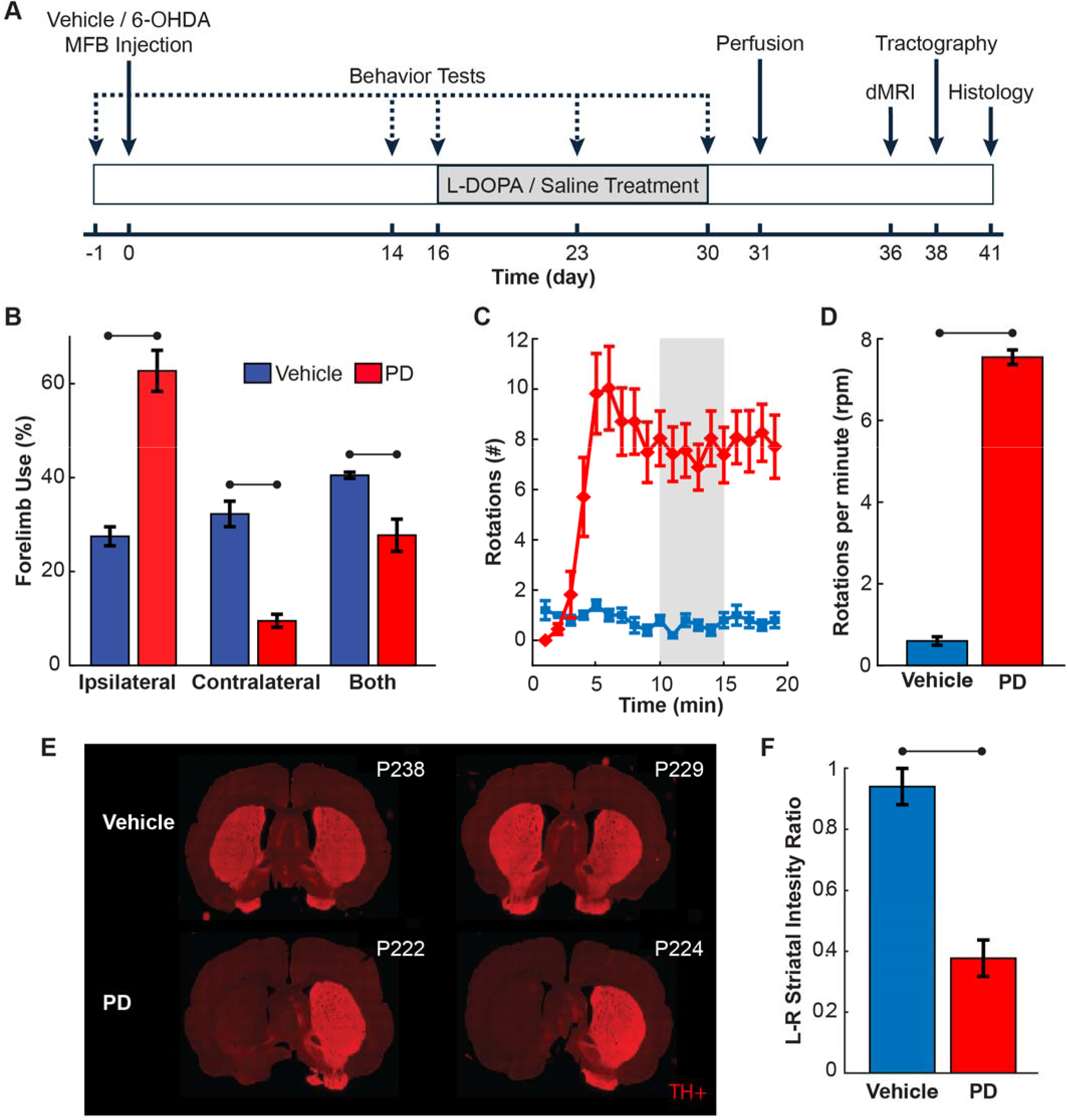
Behavioral and histological validation of the unilateral 6-OHDA PD model. Cohorts: Vehicle (blue) n = 4, PD (red) n = 22. **A** Experimental timeline. **B** Forelimb use and locomotor function evaluated from the cylinder test: percentage of ipsilateral, contralateral, and simultaneous (“both”) wall touches for Vehicle and PD rats. **C** Time course of the average contralateral rotation number over 1–20 minutes after APO injection, for Vehicle and PD rats. **D** Average rotation rate in the 10–15 minutes window after APO injection (gray window in **(C)**). **E** Representative coronal striatal slices from 2 Vehicle and 2 PD rats; bright red regions mark dopaminergic neurons staining positive for tyrosine hydroxylase (TH+). **F** Ratio of mean TH+ pixel intensity between the left versus right (i.e. ipsi-lesional vs. contra-lesional) striatum for Vehicle and PD rats. All error bars indicate mean ± 1 standard error of the mean (s.e.m.). Bars indicate significant difference at p < 0.05.

The cylinder test revealed a significant change in forelimb use between the PD and Vehicle cohorts (Fig.1B). Relative to Vehicle rats (n = 4), PD rats (n = 22) used the forelimb contralesional to their left MFB lesion far less to explore the cylinder wall (two-sample t-test, Bonferroni corrected: T₂₄ = 6.57, p = 8.55×10⁻LJ), paired with a significant increase in use of their ipsilesional forelimb (T₂₄ = −3.37, p = 0.003). Simultaneous use of both forelimbs did not differ between cohorts (T₂₄ = 1.55, p = 0.13). In conjunction, injection of APO elicited increased rotational behavior during open field locomotion consistent of dopamine-depletion in rats (i.e. PD cohort)^36–38^. PD rats began contralesional rotations about 2 minutes after injection, reached steady state roughly 7 minutes later, and persisted throughout the evaluation period (Fig 1C). In contrast, Vehicle rats rarely rotated during locomotion (Fig. 1C). Averaged over the 10–15 minutes window after APO injection (Fig. 1C, gray window), PD rats rotated significantly more than their Vehicle counterpart (Fig 1D, two-sample t-test: T_24_ = -3.20, p = 0.004).

Histology corroborated the differences in behavioral phenotypes, linking them to dopamine depletion. On coronal slices of the striatum from PD and Vehicle rats, dopaminergic neurons were identified using tyrosine hydroxylase (TH) immunohistochemistry (Fig. 1E). Striatal sections from PD rats showed 60% reduction in mean TH+ signal compared to Vehicle (PD: 0.38 ± 0.01; Vehicle: 0.94 ± 0.03, mean ± SEM; Fig. 1F; two-sample t-test: T24 = 17.31, p = 4.61e-15).

### PD-LID Model: Behavioral and Histological Confirmation

Starting on day 16 (Fig. 1A), the 22 PD rats that met the rotation criterion entered the treatment phase and were randomly assigned into 2 groups: the first group received chronic L-DOPA (6 mg/kg plus 12 mg/kg benserazide hypochloride, n = 15); the second group received saline (PD+Saline, n = 7). We scored abnormal involuntary movements (AIMs; based on amplitude of locomotor, axial, limb, and orolingual movements) across the 14-day treatment period to track the emergence of LID and its severity. Consistent with Winkler et al.^11^, AIMs peaked roughly 40–80 min after each L-DOPA injection; however, not all L-DOPA–treated rats showed this development (Fig. 2A). On the final treatment day (day 30), we classified the 15 L-DOPA–treated rats based on their integrated AIMs (Fig. 2A, B) by computing the area under the curve (AUC) of their cumulative AIMs scores. Using a fixed threshold, we defined an L-DOPA–induced dyskinesia group (LID, score > 16; n = 8) and a non-dyskinetic group (nLID, score ≤ 16; n = 7). On day 30 (Fig 2B), one-way ANOVA showed a significant effect of group on AUC of total AIMs scores (F(2,19) = 93.63, p < 0.001), with Tukey-Kramer post hoc tests showing that LID animals had significantly higher AUC than both NLID (p < 0.001) and PD+Saline animals (p < 0.001), whereas NLID and PD+Saline did not differ (p = 0.85). We also examined the rate of rotations following APO injection and TH+ staining for the PD+Saline, nLID and LID groups to verify they received similar 6-OHDA lesions. Corrected, pairwise, two-sample t-tests found no significant differences in the rate of APO rotation during the 10 - 15-minute window among the animals assigned to the LID, nLID, and PD+Saline groups (LID vs. nLID: t_l3_= −1.4720, *p* = 0.494446, LID vs. PD+Saline: t_l3_= −0.9193, *p*= 1.000000, and nLID vs. PD+Saline: t_l2_= 0.8159, *p* = 1.000000). Furthermore, examination of TH+ stained striatal sections from the LID, nLID and PD+Saline groups revealed no differences in the degree of DA depletion between the groups (Fig 2C; Tukey-Kramer post hoc (ANOVA) LID vs nLID, p=0.17; LID vs PD+Saline, p=0.99; nLID vs PD+Saline, p=0.22). See supplementary figure S1 for representative stained sections.

**Fig. 2.**
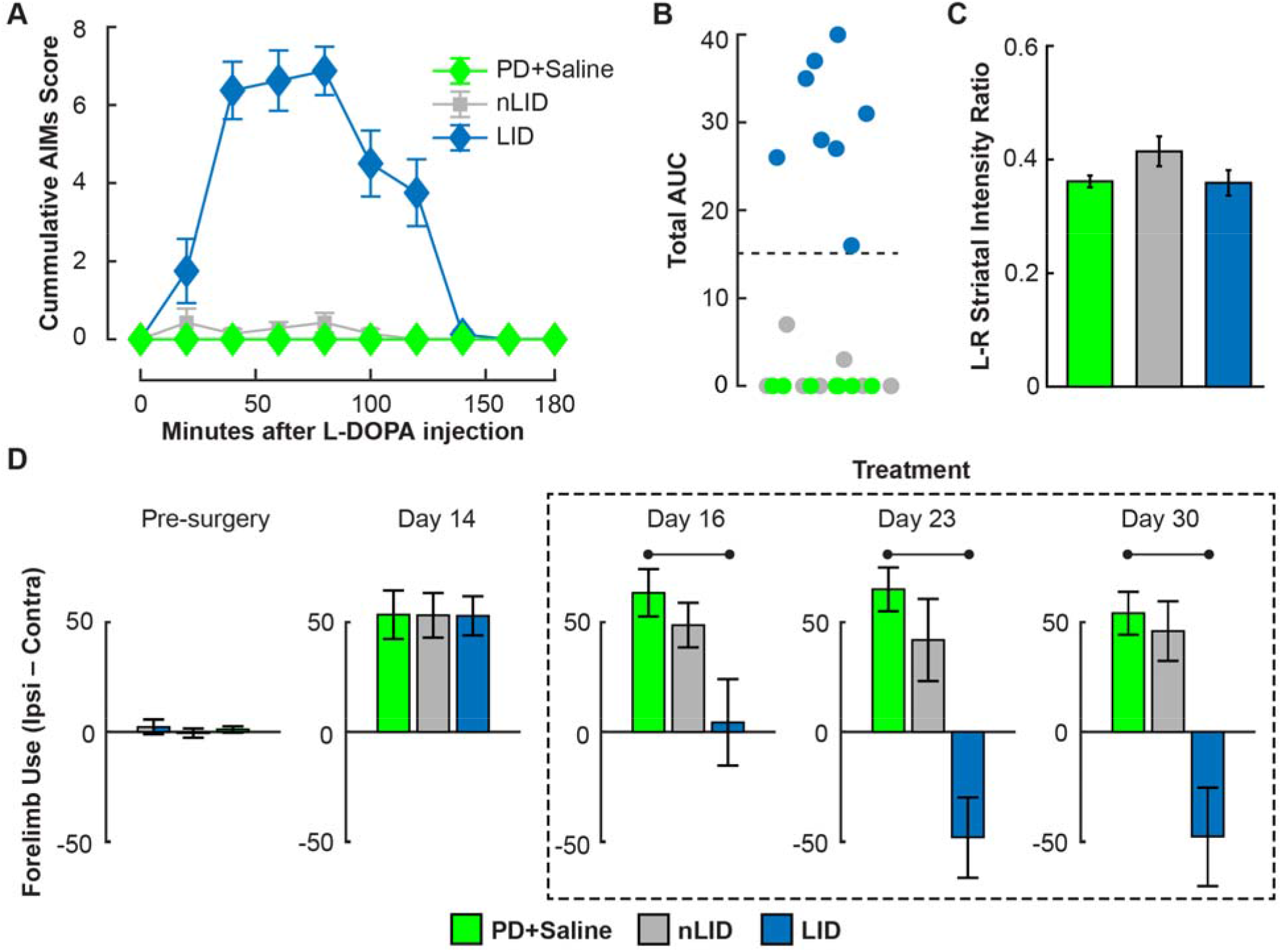
Dyskinesia development and contralateral forelimb-use reversion during chronic L-DOPA treatment. Cohorts: PD+saline (green), n = 7; nLID (gray), n = 7; LID (blue), n = 8. **A** On day 30, mean AIMs scores across 9 post-injection time windows, where the nLID and LID cohorts were injected with L-DOPA and the PD+Saline cohort received only saline. **B** Integrated AIMs score (area under the curve (AUC) of the AIMs-versus-time graphs in **(A)**) for each animal; a threshold of 16 segregated LID vs. nLID rats from the overall cohort of all PD rats who received L-DOPA treatment. **C** Ratio of mean TH+ pixel intensity between the left versus right (i.e. ipsi-lesional vs. contra-lesional) striatum of the 3 cohorts. **D** Forelimb-use asymmetry from the cylinder test (percent ipsilateral minus percent contralateral wall touches) at presurgery baseline, day 14, and treatment days 16, 23, and 30. All error bars indicate mean ± 1 s.e.m.

During the 40–80-minute window post-injection when AIMs peaked, we further tested for forelimb-use asymmetry among the cohorts using the cylinder test (Fig. 2D). Substantial asymmetry was seen in all PD cohorts (PD+Saline, nLID, LID) relative to the Vehicle cohort (data not shown); though, ANOVA found no difference between the PD+Saline, nLID and LID cohorts both at baseline (Fig. 2D, Pre-surgery) and prior to treatment (Fig, 2D, Day 14). Subsequently, symmetry was partly restored in the LID cohort on day 16 after a single L-DOPA dose (i.e. reduction in limb-use asymmetry compared to PD+Saline; ANOVA - F₂ = 4.12, p = 0.032 with post hoc Dunnett’s test – p < 0.05). Compared to PD+Saline, limb use in LID rats continued to be perturbed on Day 23 (ANOVA - F₂ = 13.74, p = 0.0002 with post-hoc Dunnett’s test – p < 0.05) and 30 (ANOVA - F₂ = 11.67, p = 0.0005 with post-hoc Dunnett’s test – p < 0.05). Here, chronic L-DOPA treatment precipitatied an inversion in limb use (negative asymmetry, i.e. increased use of contralateral compared to ipsilateral limb) in LID rats. We attribute the negative asymmetry to the emergence of dyskinetic movements. No difference in limb use between the nLID and PD+Saline groups was observed on any treatment day.

### PD Model: Tractography Results

First, we examined the structural changes underlying the PD phenotype. We used correlational tractography - a connectometry method that isolates fiber segments whose quantitative anisotropy (QA) correlates with a variable of interest - to compare 6-OHDA lesioned (PD+Saline, n = 7) versus Vehicle (n = 4) rats treated with saline, replicating our previous findings^19^. Here, PD+Saline and Vehicle were denoted categorically as 1 and 0, respectively. The analysis found fiber tracts throughout the brain whose QA was positively (red) and negatively (blue) correlated with 6-OHDA lesioning (Fig. 3A). Importantly, connectometry corroborated histological dopamine depletion; QA was reduced along the nigrostriatal fibers connecting the left substantia nigra (SN) and striatum in the PD cohort, confirming a successful 6-OHDA lesion of the left MFB (Fig. 3A, blue).

**Fig. 3.**
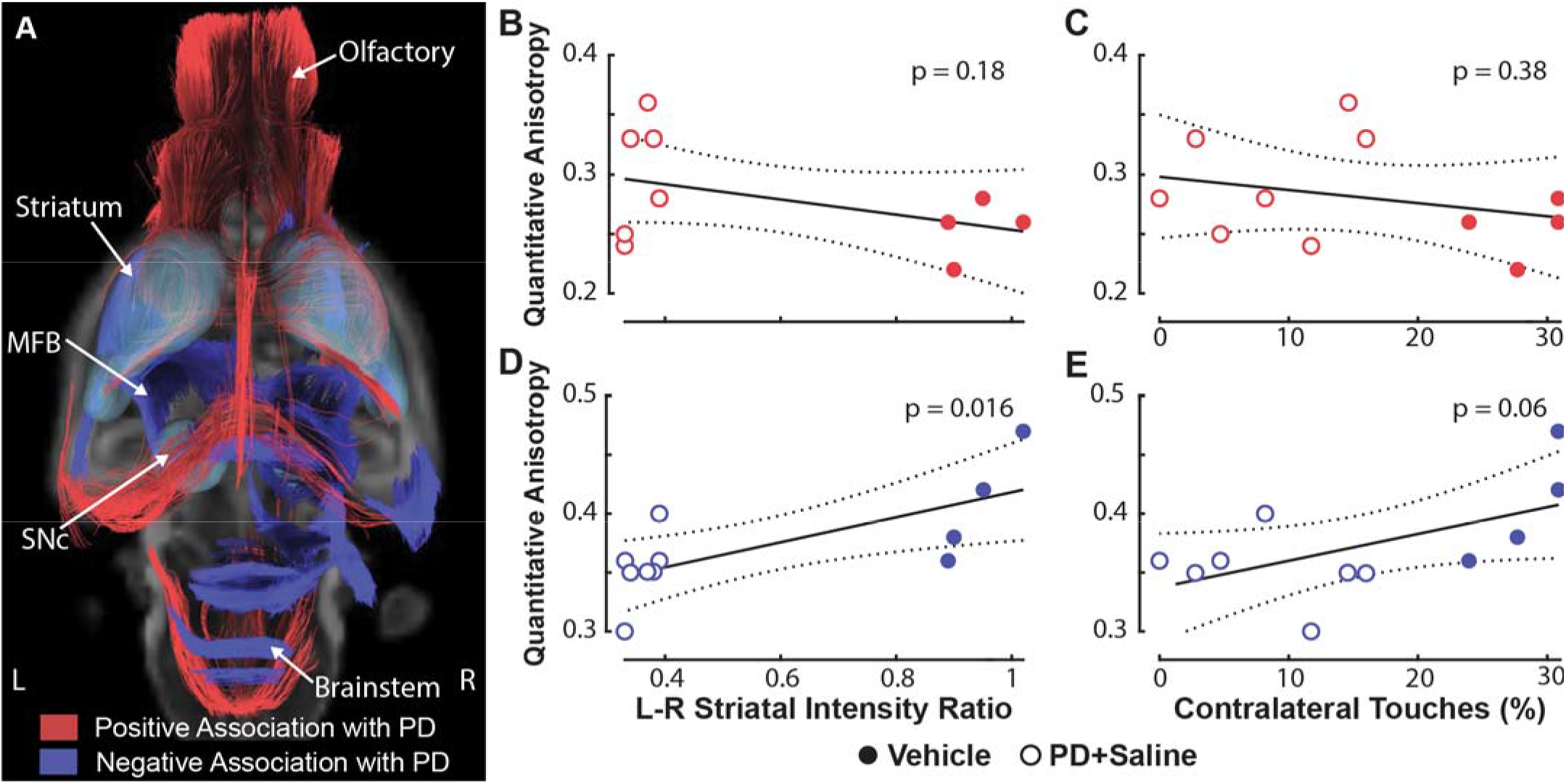
Correlational tractography describing changes in brain structure associated with unilateral (left hemisphere) 6-OHDA lesion in the MFB (PD Model). Correlational tractography comparing the PD+Saline (n=7) versus the Vehicle cohort (n =4). **A** Overlaid statistical maps of brain fiber tracts positively (red) or negatively (blue) correlated with the PD cohort (FDR < 0.05); teal volumes represent the striatum and substantia nigra. **B–E** Regression of mean QA, aggregated from positively correlated fibers **(B, C)** or negatively correlated fibers **(D, E)**, against the left-to-right striatal ratio of mean TH+ intensity **(B, D)** or the number of wall touches contralateral to the left MFB lesion **(C, E)**. Solid lines, linear fit; dotted lines, 95% confidence intervals.

QA changes extended beyond the nigrostriatal pathway, widespread across the brain. Within the lesioned, left hemisphere, fibers negatively associated with PD (Figure 3A, Blue Fibers; i.e. fibers with reduced QA in PD+Saline rats relative to the Vehicle rats) were found between the substantia nigra and striatum, striatum and piriform cortex, and striatum and ventral thalamus (Fig. 3A, blue fibers). Negatively correlated fibers were not just limited to the lesioned hemisphere; some crossed the midline to the non-lesioned side, connecting with the right ventral posteromedial nucleus (VPM) and the right ventral posterolateral nucleus (VPL) of the thalamus. Four clusters were entirely contained within the non-lesioned (right) hemisphere: (1) a posterior cluster near the right substantia nigra pars compacta (SNc) putatively connecting the periaqueductal (PAG), superior colliculus (SC), and inferior colliculus (IC) to the SNc and ventral thalamus; (2) another posterior cluster containing the cerebellar fiber tract (cbf), ventral nucleus of the lateral lemniscus, and corticospinal tract; (3) an anterior cluster joining the striatum to frontal cortex, which includes the olfactory bulb and the inferior frontal cortex (IFC, covering the olfactory, insular, piriform, and entorhinal areas); and, (4) an inferior cluster of hippocampus and amygdala connections. Like the negatively correlated fibers, positively correlated fibers (i.e. those whose QA increased in the saline-treated PD rats relative to Vehicle rats) were also widespread bilaterally, densest through the brainstem, hippocampus, striatum, and corpus callosum (Fig. 3A).

In addition, to test whether imaging changes reflected the degree of lesioning or the behavioral phenotype, we regressed mean QA against striatal dopamine depletion (TH+ intensity ratio) and forelimb asymmetry (contralateral wall touches). QA in the positively correlated fibers related to neither dopamine depletion (Fig. 3B; two-sided linear regression, t₉ = −1.47, p = 0.18) nor forelimb use (Fig. 3C; t₉ = −0.92, p = 0.38). In contrast, QA in the negatively correlated fibers was significantly correlated with the degree of dopamine depletion (Fig. 3D; t₉ = 2.96, p = 0.016) and trended toward significance with forelimb use (Fig. 3E; t₉ = 2.12, p = 0.06).

### PD-LID Model: Tractography Results

Chronic L-DOPA treatment in the PD cohort led to the development of LID in only 53% of rats (8 of 15 that received L-DOPA). Using correlational tractography, we asked what structural changes accompany LID expression. Correlating QA with the integrated AIMs score among the LID, nLID, and PD+Saline groups – with the PD+Saline cohort included to control for potential treatment effects, we identified broad, bilateral fiber networks that either positively (green) or negatively (yellow) correlated with dyskinesia severity (Fig. 4A). Positively correlated fibers were most prominent in connections between the left motor cortex (MC) and left striatum, nucleus accumbens and olfactory system. Clusters of positively correlated fibers were also found on the contralateral (right) hemisphere, including the striatum, hypothalamus, amygdala, nucleus accumbens, prefrontal cortex, piriform area, and olfactory system. Negatively correlated fibers connected the left striatum, SN, cerebellum, entorhinal (ENT) and piriform cortices, hippocampus, brainstem, corpus callosum, and PAG. Negatively correlated fibers had homologous contralateral (right) hemisphere involvement that additionally included the dorsal and ventral pallidum, VPM, VPL, reticular thalamic nucleus, and hypothalamus. Notably, fiber networks with either positive or negative association with AIMs extended well beyond canonical basal-ganglia motor circuitry, similar to network changes with the development of PD (Fig. 4A vs. Fig. 3A).

**Fig. 4.**
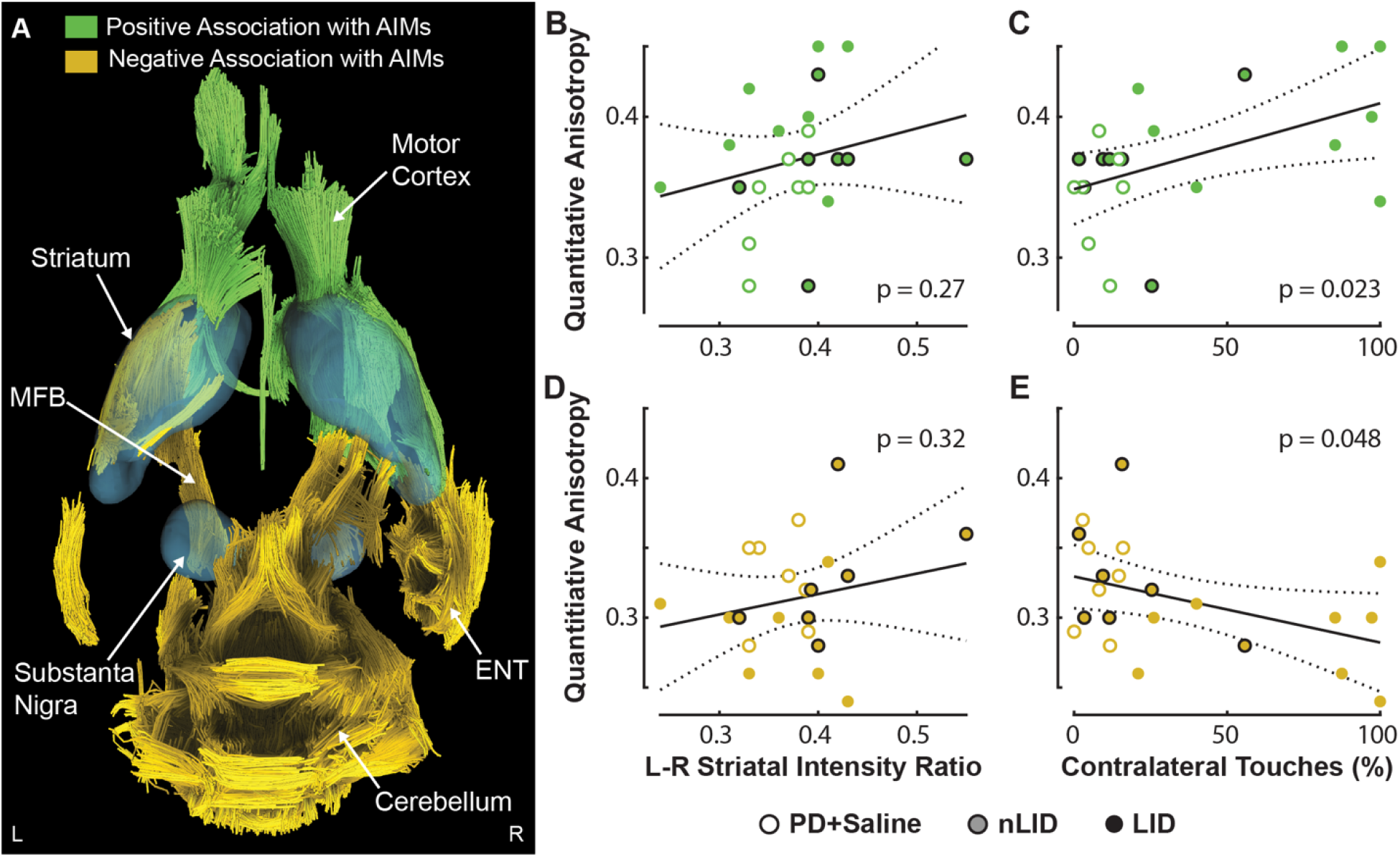
Correlational tractography describing changes in brain structure associated with AIMs following unilateral (left hemisphere) 6-OHDA lesion (PD-LID Model). **A** Overlaid statistical maps of brain fiber tracts positively (green) or negatively (yellow) correlated with the AIMs scoring (FDR < 0.05); teal volumes represent the striatum and substantia nigra. **B–E** Two-sided linear regression of mean QA, aggregated from positively correlated fibers **(B, C)** or negatively correlated fibers **(D, E)**, against the left-to-right striatal ratio of mean TH+ intensity **(B, D)** or the number of contralateral wall touches **(C, E)**. Solid lines, linear fit; dotted lines, 95% confidence intervals.

A simple linear regression analysis was again conducted to determine the degree to which the QA fibers positively and negatively correlated with AIMs were predicted by forelimb use and dopamine depletions. QA in both the positively and negatively correlated AIMs fibers scaled with forelimb use (Fig. 4C; t₂₀ = 2.47, P = 0.023; Fig. 4E; t₂₀ = −2.11, P = 0.048) but showed no relationship to dopamine depletion (Fig. 4B; t₂₀ = 1.13, P = 0.27; Fig. 4D; t₂₀ = 1.01, P = 0.32).

Since the correlational tractography of both the PD Model (Fig. 3A) and then, the PD-LID Model (Fig. 4A) implicated the SN–striatal fibers on the lesioned hemisphere, we explored whether they were the same tracts in both comparisons. We superimposed the statistical maps for both analyses with a focus on the motor cortex, striatum, and substantia nigra within the lesioned hemisphere showing that fibers were spatially segregated and distinct (Fig. 5A). Fibers negatively associated with the development of PD Model (blue) ran between the SNc and striatum; these negatively associated fibers likely represent degenerating dopaminergic projections. In contrast, fibers associated with dyskinesia (PD-LID Model) ran between the motor cortex and striatum (green), and striatum and substantia nigra pars reticulara (SNr, yellow). The dyskinesia tracts thus occupied nigrostriatal territory distinct from the lesioned dopaminergic pathway, with the striatal-SNr component potentially reflecting direct-pathway spiny projection neurons (dSPNs).

**Fig. 5.**
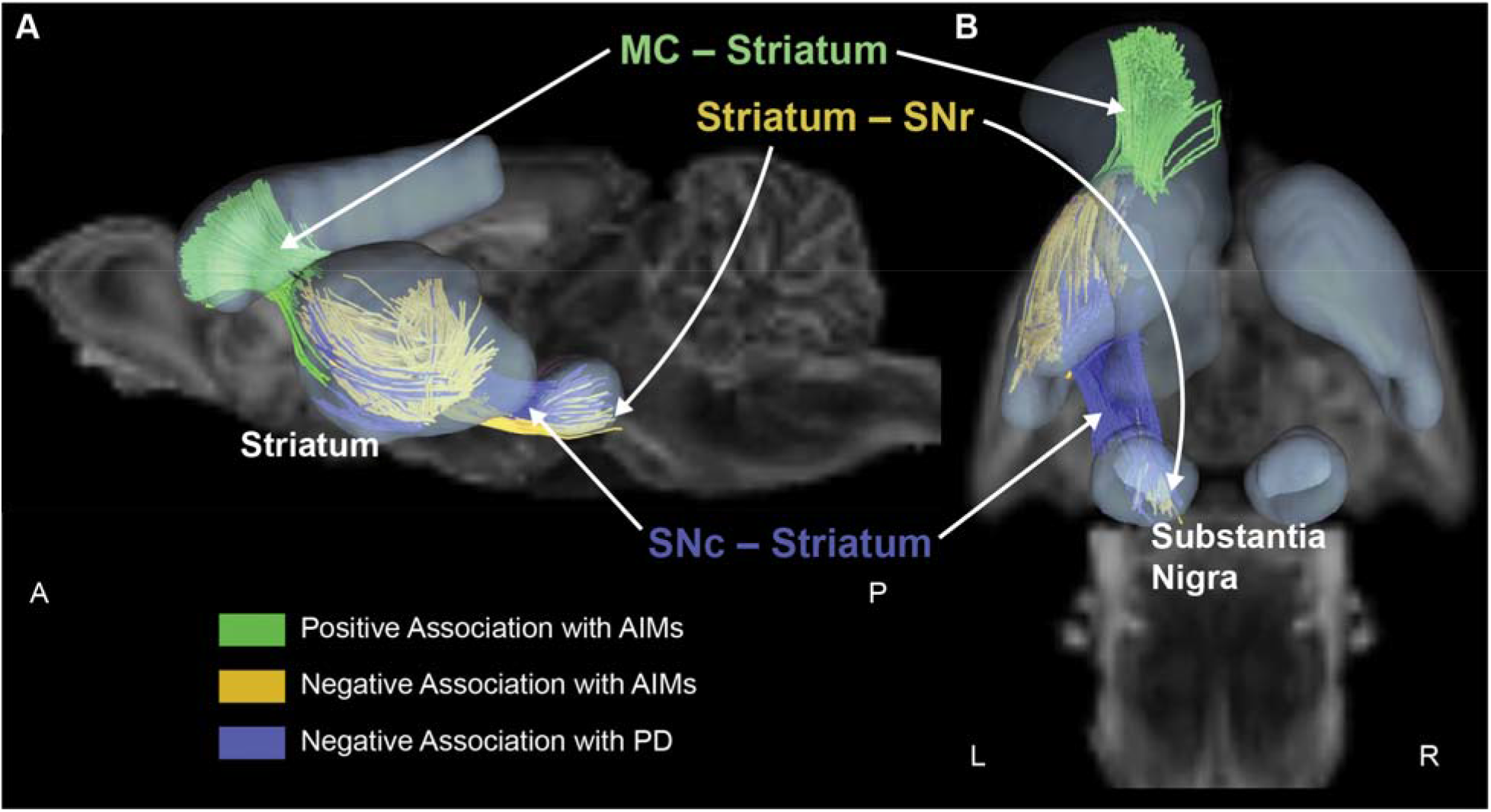
Structural changes associated with PD-LID Model versus the 6-OHDA lesion (PD Model) occupy anatomically distinct nigrostriatal pathways. Correlational tractography of integrated AIMs across all L-DOPA and saline-treated PD rats. **A–B** Brain statistical maps overlaid with sagittal **(A)** and axial **(B)** views of the lesioned hemisphere (FDR < 0.05). Fibers negatively correlated with the PD phenotype (blue; SNc–striatum connections, marking lesioned DA neurons) are spatially segregated from AIMs-correlated fibers (positively correlated: green, motor cortex–striatum; negatively correlated: yellow, substantia nigra pars reticulara (SNr)–striatum). Teal volumes represent the striatum, substantia nigra (SN), and motor cortex (MC).

Regressions of striatal dopamine depletion (TH+ intensity ratio) or forelimb asymmetry (contralateral wall touches on Day 30) with mean QA of fibers in these regions of interest reinforced this separation (Fig. 6A–E). Among the negatively associated with PD-Model fibers, the within-striatal compartment was unrelated to either measure (Fig. 6A; DA depletion, *t*₂₀ = 2.02, *p* = 0.07; forelimb use, *t*₂₀ = 1.67, *p* = 0.13), whereas the SNc–striatal projection scaled with both (Fig. 5B; DA depletion, *t*₂₀ = 3.60, *P* = 0.006; forelimb use, *t*₂₀ = 2.50, *p* = 0.03). Among the AIMs fibers, the positively correlated corticostriatal (MC–striatum) tract related to neither measure (Fig. 6C; DA depletion, *t*₂₀ = −0.26, *p* = 0.8; forelimb use, *t*₂₀ = 0.74, *p* = 0.5). In contrast, the negatively correlated AIMs fibers displayed differences based on their anatomical location. Fibers within the striatum scaled with dopamine depletion but not forelimb use (Fig. 6D; DA depletion, *t*₂₀ = 2.60, *p* = 0.016; forelimb use, *t*₂₀ = −0.39, *p* = 0.7), whereas the striatal–SNr 319 projection scaled with forelimb use but not depletion (Fig. 6E; forelimb use, *t*₂₀ = −2.20, *p* = 0.039; DA depletion, *t*₂₀ = 0.84, *p* = 0.41).

**Fig. 6.**
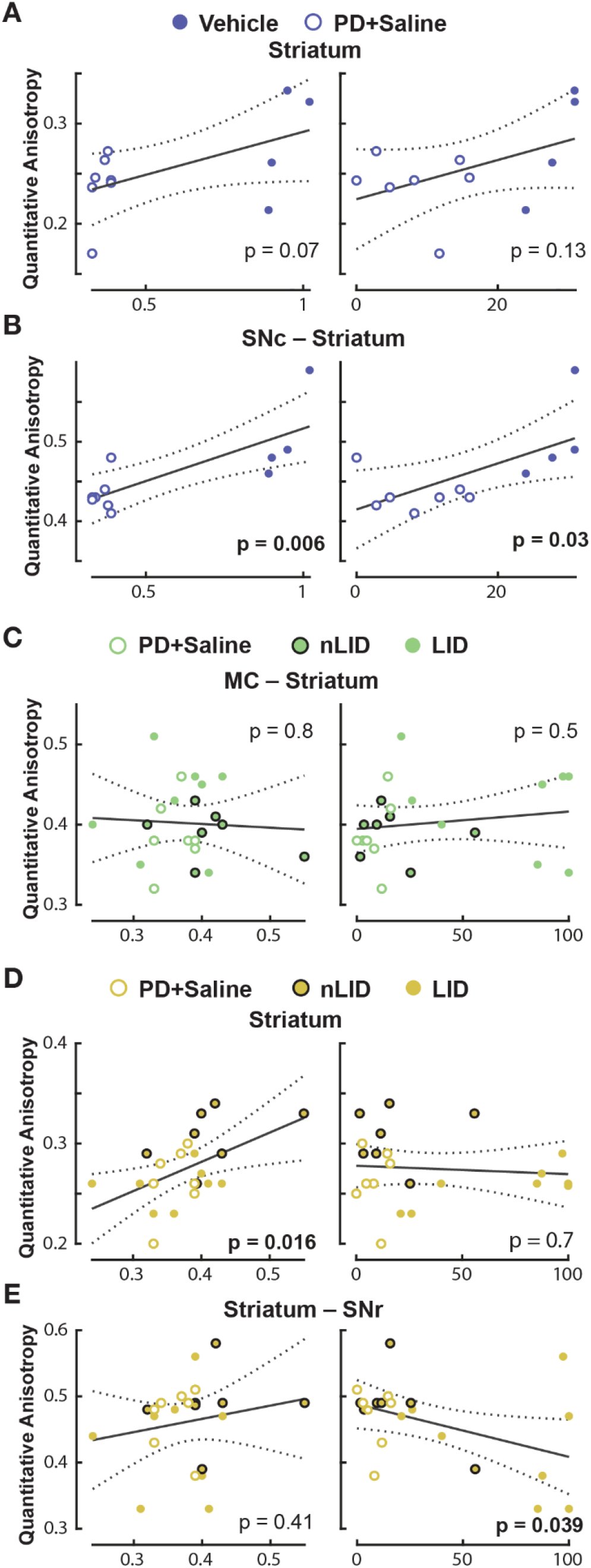
Linear regression of fiber tracts on lesion side in PD and PD-LID Models with TH+ intensity and contralateral wall touches. A–E Linear regression of mean QA against the left-to-right striatal ratio of mean TH+ intensity or the number of contralateral wall touches on Day 30, for each fiber population of interest: PD-Model, negatively correlated, within striatum (A); PD-Model, negatively correlated, SNc–striatum (B); AIMs, positively correlated, MC–striatum (C); AIMs, negatively correlated, within striatum (D); AIMs, negatively correlated, SNr–striatum (E). Solid lines, linear fit; dotted lines, 95% confidence intervals.

## Discussion

PD is commonly identified by symptoms such as resting tremor, bradykinesia, and gait disturbances. These symptoms are associated with DA deficiency and often respond positively to dopamine replacement therapy^39^. The primary pharmacological treatment for PD is L-DOPA, a dopaminergic precursor that serves as an external source of dopamine and can alleviate many of the symptoms of PD^39,40^. However, long-term treatment with L-DOPA can lead to complications, including motor oscillations and LID^7,9,41^. Many have previously explored common factors related to the development of LID (i.e. medication dose^9,33^, PD severity^33^, age of onset^42^, and its maladaptive effects on neural circuits interacting with basal ganglia structures directly impacted by PD pathophysiology (SPN changes^43^, GABAergic^44^, serotonergic^39,45–48)^ and pre/post synaptic glutamatergic^49^. A growing consensus points toward a comprehensive neurocircuit based perspective on the mechanisms of LID implicating changes in canonical nigrostriatal circuits as well as connected regions extending well beyond the borders of the dopaminergic system^50^. Here, we provide new evidence using dMRI supporting this perspective. Specifically, we show that LID severity, measured using AIMs, is strongly correlated with changes in brain microstructure (i.e. QA) in separable fiber pathways linking nodes within basal ganglia, frontal cortex, thalamus and cerebellum. Importantly, changes in brain microstructure related to LID were distinct from those identified in our PD model (identified previously^19^ and confirmed here). These results support the notion that LID emergence underlies broad systemic dysfunction stemming from maladaptive changes in nigrostriatal circuits caused by long-term L-DOPA therapy, and raise the possibility that LID-like symptoms may extend to other non-motor domains.

### Behavioral and dMRI alterations associated with LID

Our AIMs results align with previous findings in the well-established unilateral 6-OHDA rat model PD^11,51^. We found that a therapeutic dose of L-DOPA^11^ (6 mg/kg) led to short-term improvements in L-DOPA responsive rats (Fig. 2D; Day 16), but treatment effects diminished, and severe LID emerged after a few days of treatment (Fig. 2D, Day 23 and 30). Interestingly, 53% of PD rats that recieved treatment were L-DOPA responsive and developed LID (n=8), while the remaining did not (n=7). This was curious given that all animals were lesioned similarly (Fig 2). Putterman et al.^33^ showed similar variability in the development of LID responses and demonstrated how L-DOPA dose and level of striatal DA depletion drive LID. It is important to note that there has been limited focus on the non-dyskinetic animals that received chronic L-DOPA treatment. For instance, Picconi et al.^52^ included non-dyskinetic animals in their study and reported improvements with a therapeutic dose of L-DOPA (10 mg/kg) in the non-dyskinetic group (nLID) compared to the dyskinetic group (LID). In their study, the positive pharmacological effects were evaluated using the rotarod and limb-use asymmetry tests. However, our behavioral assessments using the cylinder test did not indicate any motor improvement or LID development in the nLID group. Future work should consider using a broader battery of behavioral measures to assess treatment efficacy as the activation effect of L-DOPA may be highly task dependent in some rats.

The variability in the LID phenotype for L-DOPA treated rats provided a framework to explore how brainwide neural circuits are impacted by long-term DRT. Using correlational tractography, we found that alterations in brain microstructure associated with LID severity were not limited to cortico-basal ganglia circuits in the lesioned hemisphere, but rather extended into other non-motor and integrative areas, including cerebellum, prefrontal cortex and limbic regions. Notably, the number and density of fibers negatively correlated with LID severity was asymmetric in the striatum and thalamus showing increased involvement of structures in the non-lesioned compared to the lesioned hemisphere. Moreover, the observed structural changes were not strongly related to the extent of DA depletion or limb use asymmetry, suggesting that LID-associated circuit alterations may reflect broader network-level adaptations that are not explained solely by dopamine loss or the motor phenotype, alone (Fig. 4). Importantly, the nature and effect of LID associated structural changes may vary based on the proximity of impacted networks to DA depleted regions, the relative importance of DA to local computations or how brainwide changes in glial sensitization impact input/output relationships for local neural computations^53^. Transcriptomics studies have revealed a host of gene expression changes following 6-OHDA, L-DOPA administration, and/or LID, shedding light on the potential cellular and molecular correlates of underlying plasticity-related mechanisms. For example, a 14-day course of L-DOPA, delivered to a mouse model of inducible, progressive dopaminergic degeneration, resulted in widespread and region-specific changes in gene expression in both motor and non-motor brain areas. Notable upregulation of activity-related genes (*Fos*, *Egr1*, and *Junb*) was observed in dorsal striatal and dorsal cortical (layers II/III and V) areas^54^, and L-DOPA modulated the expression of plasticity-associated and glutamatergic markers in piriform cortex and dorsal striatum, respectively. A study employing spatial multi-omics assays^55^ assessed the relationship between lipid remodeling and spatial proteomics profiles in 6-OHDA lesioned mice, revealing region-specific changes in synaptic markers and indicators of depressed cellular metabolism.

While the experimental variables vary widely across available literature (disease models, species, L-DOPA dosing, molecular assays used), these studies coalesce around observations of regionally-specific plasticity-related, metabolic, and activity-dependent changes in gene expression at the cellular level: effects that have the potential to drive broader, circuit-level restructuring. Further investigation of the cellular and molecular correlates of the LID-associated microstructural changes revealed here is a subject of future work.

### LID related structural changes in DA depleted cortico-basal ganglia circuits

LID is most often discussed in the context of its impact on the molecular, cellular, and circuit function within the DA depleted cortico-basal ganglia circuit. As expected our tractography results demonstrated robust, distinct changes in brain microstructure both positively and negatively correlated with LID severity in the lesioned hemisphere. One prominent feature was the large group of fibers connecting the motor cortex and striatum that were positively associated with LID severity (Fig.5A, green fibers). Interestingly, these fibers appeared to be strongly responsive to LID severity only given their average QA was not correlated with the degree of DA depletion (Striatal Intensity Ratio) or motor behavior (Fig. 6C). Indeed, these results are consistent with data showing chronic L-DOPA treatment alters corticostriatal plasticity^26,52,56–58^. Picconi et al.^52^ demonstrated that long term L-DOPA treatment at a therapeutic dose causes abnormal plasticity of corticostriatal connections associated with LID development. Additionally, MRI studies^22,59,60^ show that motor cortex is hyperactive in LID. We suggest that such cortico-striatal hyperactivity may lead to increased myelination and thereby more anisotropic diffusion (i.e. increased QA) in animals with more severe LID. Indeed, recent work has demonstrated alterations in corticostriatal inputs onto SPNs was linked to LID^61–65^.

In addition to alterations in the cortical input to the basal ganglia, we found that the microstructure within separate pathways within the striatum itself are negatively correlated with LID severity and DA depletion (Fig 5), respectively. One possible explanation for reduced striatal QA (negatively correlated fibers with AIMs) is morphological changes in striatal SPNs. In a unilateral 6-OHDA mouse model, Fieblinger et al., 2014 showed that DA depleted and LID mice both have reduced dendritic arborization compared to healthy controls, and that those with LID have lower number of dendritic spines for dSPNs. Overall QA in these fibers was only related to the degree of DA depletion (Striatal Intensity Ratio) and not motor behavior (Fig. 6A and D) indicating a role of the underlying pathophysiology on these striatal responses. Changes in dendrite arborization and density are known to be strongly, linearly correlated to dMRI properties^66^. We find support for the finding of separable striatal fibers correlated with LID and PD in recent work by Ryan et. al.^61^. They showed that mice with LID possess a population of dSPNs whose pattern of excitatory input, DA receptor expression and excitability are distinct from those with PD alone. Distinct gene expression profiles have been observed in rats developing AIMs versus those that did not (despite evidence of motor improvement)^67^. In rats with dyskinesia, plasticity-related gene expression (i.e., genes related to synaptic plasticity, calcium homeostasis and signaling) was upregulated. D1 receptor expression was increased in dyskinetic rats while D2 receptor expression remained unchanged. A more recent study employing single-nucleus RNA sequencing (snRNA-seq) revealed gene expression changes in D1-expressing dSPNs following LID in male 6-OHDA-lesioned mice, particularly related to genes governing synaptic remodeling and plasticity^68^.

Like the striatum, we found separable fiber tracts negatively associated with DA depletion and LID severity connecting the striatum and SN. Replicating our previous work^19^, fibers negatively associated with disease state (PD-Model; Fig 3A and 5) connected the SNc and striatum and were significantly correlated to both DA depletion and motor behavior (Fig 6B). Given the location of our 6-OHDA injection in the MFB and the confirmed DA loss in the striatum (via TH+ histology; Fig 1E), we interpret these findings as the degration of dopaminergic neurons projectng from the SNc to striatum. This is consistent with reduction in diffusion anisotropy (i.e. QA) being linked to axonal damage^69^. Conversly, LID severity was linked with a distinct bundle of fibers connecting the striatum and SNr (Fig 4 and 5) a pathway canonically associated with striatal output via the direct pathway^70^. The average QA extracted was correlated with limb use asymmetry measured on day 30, but not the degree of DA depletion (Fig. 6E). The relationship between motor behavior and QA in Striatum-SNr fibers likely reflects dyskinesia severity as motor behavior is strongly impacted by dyskinesia. Support for this assertion is found in the work of Fujita and colleagues^71^. Using a similar unilateral 6-OHDA model, they showed that rats who developed LID after long-term DRT possessed significantly reduced dendritic spine density in direct pathway neurons labeled by retrograde tracer injections. Such cellular changes are consistent with reduced QA as a less structured environment promotes greater isotropic diffusion.

### Cerebellar involvement in LID

The role of the cerebellum in the development and progression of LID remains controversial. Emergin evidence suggests that compensatory responses in the cerebellum may contribute to LID^72,73^ and that the cerebellum is a viable target for LID therapeutic interventions^74,75^. For example, Koch et al.^75^ used transcranial magnetic stimulation (TMS) on the lateral cerebellum in dyskinetic patients with Parkinson’s disease, which resulted in a reduction of LID symptoms. Additionally, Coutant et al.^74^ employed optogenetic stimulation of purkinjie cells in a freely moving mouse model of LID, leading to improvements in symptoms. A large portion of the statistical map negatively associated with AIMs score was found in the cerebellar cortex (see Fig.4A) demonstrating broad changes in tissue microstructure correlated to LID severity. Given the known structural and functional connectivity between the cerebellum and the basal ganglia^76,77^ it is not suprising that cerebellar function contributes to LID development. Future work is necessary to better understand the cellular/molecular mechanisms that lead to our dMRI finding. While direct evidence is limited, it is possible that the unnatural, cyclic administration of L-DOPA promotes neuroinflamation and glial proliferation^78^ within the cerebellum disrupting the firing dynamics of cerebellar output nuclei^73^. Changes in cerebellar output are likely due to abberant firing of Purkinje cells which have been tied to LID via changes in sigma receptor density^79^. Here, we expect that alterations in microglia morphology and astrocyte proliferation would contribute to the inverse relationship between QA and LID severity, specifically reduced diffusion anisotropy with increased dyskinesia severity.

### LID implications for non-motor domains

In addition to canonical regions on the brain known to contribute to the motor symptoms of PD and LID, we also found many regions associated with AIMs severity in regions putatively linked to non-motor functions like cognitive function/dementia and pain. In both the lesioned and nonlesioned hemispheres, correlation tractography found the negatively correlated fibers associated with AIMs within many brain regions including the entorhinal cortex (ENT), the piriform cortex, the hippocampus and and the periaqueductal gray matter (see Fig. 4A). Notably, atrophy of the entorhinal cortex in PD can serve as a marker to distinguish between patients with dementia and those with normal cognition^80,81^. Furthermore, hippocampal atrophy is often observed in PD^82,83^. Notably, Yang, et al.^84^ demonstrated that rats with unilateral 6-OHDA lesions exhibit a reduction in the cytomembrane of hippocampal neurons, which is associated with increased anxiety compared to control animals. In a similar model, Kim et al.^85^ demonstrated changes in the dendritic complexity of hippocampal cornu ammonis 1 (CA1) neurons on the lesioned side. Like memory and cognition, pain-related signaling is often impacted in in patients with late-stage PD implicating the periaqueductal gray matter^86^. In unilateral 6-OHDA models, pain-related symptoms associated with PD and L-DOPA therapy have garnered considerable attention. For example, unilateral 6-OHDA lesion caused bilateral reductions in nociceptive thresholds when exposed to heat, mechanical, and cold stimuli^87^, and that normal thresholds are restored by L-DOPA treatement^88^. Future work should address if LID-like symptoms are present in non-motor domains.

## Limitations

Here, we present data supporting comprehensive neurocircuit based perspective on LID implicating brainwide changes in microstructure as a consequence of its pathology and subsequent compensatory responses. There are several limitations regarding our experimental design and techniques that should be considered when interpreting the results of this study. First, dMRI is - by nature - an indirect measure of the changes caused by LID with the underlying molecular and cellular mechanisms being veiled. We used histological evidence to validate that the negative association between QA within PD+Saline and Vehicle cohorts was caused by degradation of the MFB precipitated by 6-OHDA. However, the link between some QA associations, both negative and positive, and underlying physiology are subject to speculation as the cause of changes in measures of anisotropy are multifactorial. Future work is necessary to specifically link changes in QA associated with LID severity to variations in histological markers or transcription factors. Second, we used changes in striatal TH intensity as a marker for loss of DA neurons in the nigrostriatal system. While these data reflect the expected changes caused by the 6-OHDA lesions in the MFB, they do not specifically quantify differential loss of SNc neurons as would be quantified by sterology. Future, work could consider counting neurons in the SNc. Third, unilateral 6-OHDA lesions result in rapid and severe loss of dopamine neurons and may not fully recapitulate the progression of clinical PD or the multi-year development of LID PD^89,90^. Furthermore, we did not apply an individualized L-DOPA dosing paradigm for the rats in this study which may have altered the progression of LID. Therefore, we must exercise caution when translating our findings to humans with PD as they all have unique comorbidities and history. Third, we chose to examine dMRI related changes in fixed, post-mortem tissue to achieve the quality imaging possible as having high resolution, high signal-to-noise ratio (SNR) images is a key factor for accurate tractography^91^. That said, fixation changes the diffusivity of water molecules compared to living tissue^92,93^, but adjustments of imaging parameters similar to the ones we employed (i.e. increased numbers of averages and higher B-values) compensate for changes due to these effects^91^. Fourth, it is possible that our data measuring limb-use asymmetry at days 23 and 30 were impacted by the development of LID and thus reflect the development of LID alongside changes in limb preference. This effect was likely impacted by our choice to measure AIMs during peak dose dyskinesia. Future work should consider alternate measures of motor behavior not impcted by LID or measuring before/after peak LID symptoms.

## Methods

### Animals

Fifty-four male Long Evans rats (12 weeks of age, weighing 313–403 g) were used for this study, where they were housed on a 12-hour light/dark cycle with ad libitum access to food and water. All animal procedures were approved by the Institutional Animal Care and Use Committee (IACUC) at the University of Wisconsin–Madison.

### Stereotactic Surgery

Rats were anesthetized with isoflurane in O₂ (5% induction, 1–2.5% maintenance) and placed in a stereotaxic frame. To protect noradrenergic neurons from the subsequent 6-OHDA lesioning, each rat also received desipramine hydrochloride (20 mg/kg; Sigma-Aldrich) at least 30 minutes before the stereotaxic injection^94^. We then shaved the top of the head, cleaned the skin with povidone-iodine and alcohol scrubs, incised the scalp at the midline, and retracted all tissue to expose lambda and bregma. We leveled the skull and drilled a small burr hole above the injection site (−4.3 mm AP, −1.6 mm ML). Under microscopic guidance, we confirmed contact of the needle with the dural surface and then lowered it to the MFB (−8.0 mm DV). Upon reaching the target depth, we idled the needle for 5 minutes. Throughout the surgery, we monitored heart rate, SpO_2_, and body temperature (PhysioSuite, Kent Scientific), where a heating pad was further used to maintain body temperature. Rats were initially segregated into either the 6-OHDA or the Vehicle cohort. The 6-OHDA cohort received unilateral, 6-OHDA injections into the left MFB (8 μg in 2 μl of saline of saline with 0.02% ascorbate over 8 minutes; Sigma-Aldrich). In contrast, the Vehicle cohort received injections of sterile phosphate-buffered saline 2 μl over 8 min. After injection, we idled the needle for 5 minutes before withdrawing it. Finally, the burr hole was covered with gel foam soaked in sterile saline and the skin incision closed with 5-0 Vicryl suture.

### Apomorphine Test

On day 14, we conducted an apomorphine test to confirm the PD phenotype in our rat model^36–38^. We injected the rats subcutaneously with APO (0.1 mg/kg dissolved in 0.2% ascorbic acid and prepared in 0.9% sterile saline), placed them in an open plexiglass cube (303 mm sides), and video-recorded them for 30 minutes^35,95^. In this test, rats with a sufficiently lesioned left MFB were expected to rotate more frequently and away from the lesioned left side (contralateral rotations to the right), whereas non-lesioned rats should rotate less and show no directional preference^36–38^. We quantified the number of contralateral rotations using the automated motion-tracking package DeepLabCut (see Video Tracking Analysis from Moshchin et al.^19^ for details). Here, we used a two-sample t-test to compare rotation counts between the PD and Vehicle cohorts.

### Cylinder Test

The cylinder test was applied for two purposes: 1) to validate PD model on day 14 and 2) assess the effect of L-DOPA treatment on days 16, 23, and 30. Rats were placed in a glass cylinder (160 mm diameter, 395 mm height) without prior acclimatization and video-recorded for 5 minutes. From the videos, we counted each instance in which a rat used its forelimbs to navigate the cylinder wall and classified the contact as with respect to the left MFB lesion - ipsilateral to the lesion, contralateral to the lesion, or both. We then expressed each contact type as a percentage of total touches^96^:

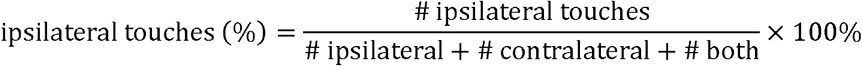

Contralateral and simultaneous (“both”) percentages were calculated analogously. We used two-sample t-tests to compare the percentage of ipsilateral, contralateral, and simultaneous touches between the Vehicle and PD cohorts, applying a Bonferroni correction to control for multiple comparisons.

### L-DOPA Treatment and the Abnormal Involuntary Movement (AIM) Rating (PD-LID Model)

Animals were allowed to recover for two weeks after 6-OHDA (PD cohort) or saline (Vehicle cohort) injection into the left MFB, after which we initiated chronic L-DOPA treatment for the PD cohort on day 16 and continued it for 14 days. We randomly segregated the PD cohort into 2 treatment arms: the first (n = 15) received daily subcutaneous injections of L-DOPA (L-3,4-dihydroxyphenylalanine, 6 mg/kg; Sigma-Aldrich) combined with benserazide hydrochloride (12 mg/kg; Sigma-Aldrich), and the second (n = 7) received daily subcutaneous saline. We held L-DOPA dosage constant throughout this experiment. After each injection, we placed rats in a glass cylinder (200 mm diameter, 400 mm height, lined with bedding to absorb waste) without prior acclimatization and video-recorded them for 3 hours. Rats were expected to develop L-DOPA induced dyskinesia (LID) to different degrees^33^. Thus, we scored the severity of LID using the Abnormal Involuntary Movement (AIM) scale during nine 1-minute monitoring windows at 20-minute intervals post-injection, as described by Winkler et al.^11^. For each window, we scored the amplitude of locomotor, axial, limb, and orolingual movements from 0 to 4, giving a maximum total of 16. Severity rating was not taken into account. We then computed each animal’s integrated AIM score (Total AUC) as the area under the AIM-score-versus-time curve (Fig. 2B).

### Perfusion

We perfused rats on day 31. Each rat was anesthetized with isoflurane (2–5%) and transcardially perfused with ice-cold PBS followed by 4% paraformaldehyde (PFA). We then extracted the brain, post-fixed it in 4% PFA for 48 hours, and transferred it to 1× PBS after washing.

### Diffusion MRI: Image Acquisition and Preprocessing

On day 36, we removed each brain from 1x PBS and placed it in a 10 mL syringe of Fluorinert (FC-40, 3M, St. Paul, MN). Syringes were grouped in batches of three and then imaged using dMRI similar to our previous work^19,97^. Prior to imaging, the orientation of the syringes within the scanner was documented photographically to preserve the identity of each sample within the grouped 3D volume. Multi-slice diffusion-weighted spin echo images were obtained with a 4.7 T Agilent MRI system fitted with a 3.5 cm-diameter quadrature volume radiofrequency (RF) coil, using the following parameters: TR = 3400 ms, TE = 24 ms, matrix = 128x128 (reconstructed to 256 x 256), field of view (FOV) = 32x32 mm, voxel size = 250 µm, section thickness = 0.35 mm (resampled to 0.25 mm isotropic in post-processing), number of excitations (NEX) = 2, and two shell acquisitions with 10 directions at b = 0 s/mm², 25 directions at b = 816 s/mm², and 50 directions at b = 2,041 s/mm² (D =12.20 ms, d = 6 ms). All imaging was performed in a temperature-controlled room at a consistent 20–21 °C.

Imaging data were converted to the NIfTI format and headers were manually checked and confirmed to match acquisition parameters (0.25 mm, 0.25 mm, 0.35 mm). To correct distortions from eddy currents, FSL eddy-correct was applied (FSL, FMRIB Software Library; Version 5.0; Oxford Centre for Functional MRI of the Brain; http://fsl.fmrib.ox.ac.uk/fsl/). Next, the imaged 3D volume aggregate was imported into DSI-Studio, resampled to isotropic (ISO) voxel size (0.25 mm in each direction), and divided into the 3 individual brains for further processing. Individual images were aligned to the Anterior Commissure-Posterior Commissure (AC-PC) orientation using the fslreorient2std tool and each image’s affine transformation was adjusted toward the standard template using the FSLeyes nudge tool. Subsequently, the sum of all transforms was applied using FSL FLIRT (FMRIB Linear Image Registration Tool) to resample the image into standard atlas template space^98^. B-table accuracy was verified using an automated tool^99^. Diffusion data were reconstructed and spin distribution functions (SDF) were computed using a model-free method, generalized q-sampling imaging (GQI, ref), with diffusion sampling length ratio set to 0.2.

### Diffusion MRI: Analysis and Tractography

Next, we used correlational tractography^32^ to assess how dopaminergic depletion modulates local connectomes, extracting fiber tracts whose diffusion metrics correlated significantly with a cohort. We estimated local connectome matrices by sampling each animal’s SDFs using the local fiber directions of the rat template^98^. Then, we assembled these local connectome matrices into two distinct databases. The first database (PD Model), comprising matrices from PD+Saline and Vehicle cohorts. Nonparametric Sperman correlation was applied to assess associations between connectome features and these groups (denoted categorically - PD+Saline = 1 and Vehicle = 0). The second database (PD+LID Model) included matrices from PD animals treated with L-DOPA and their corresponding PD+Saline controls. Here, a linear regression was first used to remove the effects of treatment type (L-DOPA or Saline) on the diffusion data, and then a non-parametric Spearman correlation evaluated the associations between the local connectome matrices and dyskinesia severity (i.e. total AUC). Here, we tracked local connectomes with positive or negative correlations using a deterministic fiber tracking algorithm^100^ following default DSI-Studio parameters (T-score threshold = 2.5). From the results, we filtered for tracks by combining topology-informed pruning (4 iterations, see Yeh et al.^101^) with a 10-voxel (2.5 mm) length threshold. Subsequently, we selected tracks with a significant association to a cohort at a false discovery rate (FDR) of 0.05, computed in DSI-Studio using the method described by Yeh, et al.^102^. A total of 4000 permutations were used to estimate the empirical and null track length histograms.

### Histology

After dMRI, aside from the imaging analysis, we processed the brain tissue to stain sections through the basal ganglia for tyrosine hydroxylase (TH). Tissue was cryoprotected at 4 °C in a 0.1 M phosphate buffer with 2% dimethyl sulfoxide (DMSO) and a graded glycerol concentration series (10% for 1 day, 15% for 4 hours, then 20% for 4 days). The tissue was frozen with dry ice and sliced coronally at 40 µm thickness on a sliding microtome. The slices were then transferred to an ethylene glycol based cryoprotectant and stored at −20 °C. For immunostaining, slices were first washed in 1x TBS. Antigen retrieval was performed at 80 °C in Tris-EDTA (pH 9.0), followed by endogenous peroxidase quenching in TBS with 3% H_2_O_2_ and 10% methanol. Slices were then washed in TBS with 0.05% Triton X-100 (TBST; Sigma-Aldrich, Burlington, MA), blocked in TBST with 5% normal goat serum for 30 minutes, and incubated overnight in TBST with 1% bovine serum albumin (BSA; Calbiochem, Millipore Sigma, St. Louis, MO) and primary antiserum (Millipore Sigma MAB318; 1:5,000). After washing in TBST, slices were incubated in Alexa Fluor 594 secondary IgG (Invitrogen A11032; 1:750) for 2 hours, rinsed with TBS, mounted on a gelatin coated slide, and cover slipped with Prolong Gold antifade and DAPI (Invitrogen, Thermo Fisher Scientific, Waltham, MA). We imaged these coronal slices on an Axio Imager Z2 microscope with ZEN 3.3 software (Carl Zeiss), using a TxRed filter cube, a 4 ms shutter exposure, and 50% LED power. To quantify striatal TH expression, we measured mean-pixel intensity of TH signal in ImageJ^103^. For each brain, we analyzed three coronal striatal slices approximately +1.00 mm anterior, +0.20 mm anterior, and -1.00 mm posterior to bregma. In each slice, we outlined the caudate putamen (CPu) in each hemisphere and extracted the region’s mean pixel intensity. Subsequently, we averaged the values for both sides of each slice and calculated the ratio, dividing the average ipsilateral value by the average contralateral value.

## Author Contributions

AJS, MM, KAL, JKT and CB conceived of and designed the experiment. AJS and WBL secured funding for the project. SO, RJS and KPC performed surgical procedures. MM, EY, RJS and JED collected the behavioral data. SO performed the histological procedures. SAH developed the dMRI acquisition protocol. MM, EY, CB and AJS analyzed and interpreted the data. MM, AJS, SO, CPL, EKP, CB and KAL drafted the manuscript. All authors contributed to refining the manuscript and approved the final version.

## Acknowledgements

Support for this research was provided by the University of Wisconsin – Madison Office of the Vice Chancellor for Research and Graduate Education, the Wisconsin Alumni Research Foundation and the Department of Neurological Surgery.

## References

1. Ren, J. Parkinson’s disease as a somato-cognitive action network disorder. Nature 651, 1030–1038 (2026).

2. Hosp, J. A. The degenerome—a novel streamline-wise approach for white matter integrity in neurodegeneration. Npj Park. Dis. 12, 139 (2026).

3. Chung, S. J. Association Between White Matter Connectivity and Early Dementia in Patients With Parkinson Disease. Neurology 98, e1846–e1856 (2022).

4. Yang, K. White matter changes in Parkinson’s disease. Npj Park. Dis. 9, 150 (2023).

5. Lee, A. & Gilbert, R. M. Epidemiology of Parkinson Disease. Neurol. Clin. 34, 955–965 (2016).

6. Fabbri, M. et al. Response of non-motor symptoms to levodopa in late-stage Parkinson’s disease: Results of a levodopa challenge test. Parkinsonism Relat. Disord. 39, 37–43 (2017).

7. Jankovic, J. & Poewe, W. Therapies in Parkinson’s disease. Curr. Opin. Neurol. 25, 433–447 (2012).

8. Fridjonsdottir, E. Mass spectrometry imaging identifies abnormally elevated brain L-DOPA levels and extrastriatal monoaminergic dysregulation in L-DOPA–induced dyskinesia. Sci. Adv. 7, eabe5948 (2021).

9. Sgroi, S., Kaelin-Lang, A. & Capper-Loup, C. Spontaneous locomotor activity and L-DOPA-induced dyskinesia are not linked in 6-OHDA parkinsonian rats. Front. Behav. Neurosci. 8, 331 (2014).

10. Guridi, J., González-Redondo, R. & Obeso, J. A. Clinical Features, Pathophysiology, and Treatment of LevodopaLJInduced Dyskinesias in Parkinson’s Disease. in Parkinson’s disease 2012.1 943159 (2012).

11. Winkler, C. L-DOPA-induced dyskinesia in the intrastriatal 6-hydroxydopamine model of parkinson’s disease: relation to motor and cellular parameters of nigrostriatal function. Neurobiol. Dis. 10, 165–186 (2002).

12. Pandey, S. & Srivanitchapoom, P. Levodopa-induced dyskinesia: clinical features, pathophysiology, and medical management. Ann. Indian Acad. Neurol. 20, 190–198 (2017).

13. Parker, J. G. Diametric neural ensemble dynamics in parkinsonian and dyskinetic states. Nature 557, 177–182 (2018).

14. Crittenden, J. R. & Graybiel, A. M. Basal Ganglia Disorders Associated with Imbalances in the Striatal Striosome and Matrix Compartments. Front. Neuroanat. 5, (2011).

15. Tang, C. C., Poston, K. L., Dhawan, V. & Eidelberg, D. Abnormalities in Metabolic Network Activity Precede the Onset of Motor Symptoms in Parkinson’s Disease. J. Neurosci. 30, 1049–1056 (2010).

16. Breit, S. Bilateral changes in neuronal activity of the basal ganglia in the unilateral 6-hydroxydopamine rat model. J. Neurosci. Res. 86, 1388–1396 (2008).

17. Ma, Y. Abnormal Metabolic Brain Networks in a Nonhuman Primate Model of Parkinsonism. J. Cereb. Blood Flow Metab. 32, 633–642 (2012).

18. Santoro, M. Mapping of catecholaminergic denervation, neurodegeneration, and inflammation in 6-OHDA-treated Parkinson’s disease mice. Npj Park. Dis. 11, 28 (2025).

19. Moshchin, M. Assessing changes in whole-brain structural connectivity in the unilateral 6-hydroxydopamine rat model of Parkinson’s disease using diffusion imaging and tractography. J. Neural Eng. 22, 046005 (2025).

20. Moshchin, M. Separable, symptom specific alterations in brain microstructure associated with early-stage Parkinson’s disease. Front. Neurosci. 20, 1741159 (2026).

21. Bastide, M. F. Immediate-early gene expression in structures outside the basal ganglia is associated to L-DOPA-induced dyskinesia. Neurobiol. Dis. 62, 179–192 (2014).

22. Rajan, R. Cortical plasticity and levodopa-induced dyskinesias in Parkinson’s disease: connecting the dots in a multicomponent network. Clin. Neurophysiol. 128, 992–999 (2017).

23. Muthuraman, M. Deep brain stimulation and L-DOPA therapy: concepts of action and clinical applications in Parkinson’s disease. Front. Neurol. 9, 711 (2018).

24. Wang, L. Altered brain structural topological properties in Parkinson’s disease with levodopa-induced dyskinesias. Parkinsonism Relat. Disord. 67, 36–41 (2019).

25. Jung, J. H. White matter connectivity networks predict levodopa-induced dyskinesia in Parkinson’s disease. J. Neurol. 269, 2948–2960 (2022).

26. Cenci, M. A. & Kumar, A. Cells, pathways, and models in dyskinesia research. Curr. Opin. Neurobiol. 84, 102833 (2024).

27. Hirano, S. Metabolic networks for assessment of therapy and diagnosis in Parkinson’s disease. Mov. Disord. 24, 725–731 (2009).

28. Lerner, R. P. Levodopa-induced abnormal involuntary movements correlate with altered permeability of the blood-brain-barrier in the basal ganglia. Sci. Rep. 7, 16005 (2017).

29. Yun, J. J. Modulatory effect of levodopa on the basal ganglia-cerebellum connectivity in Parkinson’s disease. Npj Park. Dis. 11, 115 (2025).

30. Kelly, R. M. & Strick, P. L. Cerebellar loops with motor cortex and prefrontal cortex of a nonhuman primate. J. Neurosci. 23, 8432–8444 (2003).

31. Conti, M. M. Monoamine transporter contributions to L-DOPA effects in hemi-parkinsonian rats. Neuropharmacology 110, 125–134 (2016).

32. Yeh, F.-C., Badre, D. & Verstynen, T. Connectometry: A statistical approach harnessing the analytical potential of the local connectome. NeuroImage 125, 162–171, (2016).

33. Putterman, D. B. Evaluation of levodopa dose and magnitude of dopamine depletion as risk factors for levodopa-induced dyskinesia in a rat model of Parkinson’s disease. J. Pharmacol. Exp. 277–284 (2007).

34. Eusebi, P. Risk factors of levodopa-induced dyskinesia in Parkinson’s disease: results from the PPMI cohort. Npj Park. Dis. 4, 33 (2018).

35. Carman, L. S., Gage, F. H. & Shults, C. W. Partial lesion of the substantia nigra: relation between extent of lesion and rotational behavior. Brain Res. 553, 275–283 (1991).

36. Ungerstedt, U. & Arbuthnott, G. W. Quantitative recording of rotational behavior in rats after 6-hydroxy-dopamine lesions of the nigrostriatal dopamine system. Brain Res. 24, 485–493, (1970).

37. Schwarting, R. K. W. & Huston, J. P. The unilateral 6-hydroxydopamine lesion model in behavioral brain research. Analysis of functional deficits, recovery and treatments. Prog. Neurobiol. 50, 275–331 (1996).

38. Cenci, M. A. & Lundblad, M. Utility of 6-Hydroxydopamine Lesioned Rats in the Preclinical Screening of Novel Treatments for Parkinson Disease. in Animal Models of Movement Disorders 193–208 (Elsevier, 2005). doi:10.1016/B978-012088382-0/50016-5.

39. Cenci, M. A. Presynaptic mechanisms of L-DOPA-induced dyskinesia: the findings, the debate, and the therapeutic implications. Front. Neurol. 5, 242 (2014).

40. Salat, D. & Tolosa, E. Levodopa in the treatment of Parkinson’s disease: current status and new developments. J. Park. Dis. 3, 255–269 (2013).

41. Cenci, M. A. Dyskinesia matters. Mov. Disord. 35, 392–396 (2020).

42. Nishijima, H. Effects of aging on levo-dihydroxyphenylalanine-induced dyskinesia in a rat model of Parkinson’s disease. Front. Aging Neurosci. 13, 650350 (2021).

43. Fieblinger, T. Striatonigral neurons divide into two distinct morphological-physiological phenotypes after chronic L-DOPA treatment in parkinsonian rats. Sci. Rep. 8, 10068 (2018).

44. Nishijima, H. GABA storage and release in the medial globus pallidus in L-DOPA-induced dyskinesia priming. Neurobiol. Dis. 143, 104979 (2020).

45. Zhou, F. C., Bledsoe, S. & Murphy, J. Serotonergic sprouting is induced by dopamine-lesion in substantia nigra of adult rat brain. Brain Res. 556, 108–116 (1991).

46. Gil, S. The roles of striatal serotonin and l-amino-acid decarboxylase on l-DOPA-induced dyskinesia in a hemiparkinsonian rat model. Cell. Mol. Neurobiol. 30, 817–825 (2010).

47. Rylander, D. Maladaptive plasticity of serotonin axon terminals in levodopa-induced dyskinesia. Ann. Neurol. 68, 619–628 (2010).

48. Sellnow, R. C. et al. Regulation of dopamine neurotransmission from serotonergic neurons by ectopic expression of the dopamine D2 autoreceptor blocks levodopa-induced dyskinesia. Acta Neuropathol Commun 15;7(1):8, (2019).

49. Mosharov, E. V., Borgkvist, A. & Sulzer, D. Presynaptic effects of levodopa and their possible role in dyskinesia. Mov. Disord. 30, 45–53 (2015).

50. Yang, K. Circuit mechanisms of L-DOPA-induced dyskinesia (LID). Front. Neurosci. 15, 614412 (2021).

51. Conti, M. M. Effects of prolonged selective serotonin reuptake inhibition on the development and expression of L-DOPA-induced dyskinesia in hemi-parkinsonian rats. Neuropharmacology 77, 1–8 (2014).

52. Picconi, B. Loss of bidirectional striatal synaptic plasticity in L-DOPA–induced dyskinesia. Nat. Neurosci. 6, 501–506 (2003).

53. Lima, M. N., Barbosa-Silva, M. C. & Maron-Gutierrez, T. Microglial priming in infections and its risk to neurodegenerative diseases. Front. Cell. Neurosci. 16, 878987 (2022).

54. Radlicka-Borysewska, A. L-DOPA induces spatially discrete changes in gene expression in the forebrain of mice with a progressive loss of dopaminergic neurons. Mol. Neurobiol. 62, 11466–11483 (2025).

55. Lee, S. Y. Spatial multi-omics reveals region-specific molecular signatures in a 6-OHDA model of Parkinson’s disease. Npj Park. Dis. (2026).

56. Calabresi, P. Levodopa-induced dyskinesia: a pathological form of striatal synaptic plasticity? Ann. Neurol. 47,4 Suppl 1, 60–8 (2000).

57. Calon, F. Molecular basis of levodopa-induced dyskinesias. Ann. Neurol. 47, 70 (2000).

58. Cenci, M. A. & Crossman, A. R. Animal models of l-dopa-induced dyskinesia in Parkinson’s disease. Mov. Disord. 33, 889–899 (2018).

59. Cerasa, A. Future scenarios for levodopa-induced dyskinesias in Parkinson’s disease. Front. Neurol. 6, 76 (2015).

60. Herz, D. M. R esting-state connectivity predicts levodopa-induced dyskinesias in P arkinson’s disease. Mov. Disord. 31, 521–529 (2016).

61. Ryan, M. B. Excessive firing of dyskinesia-associated striatal direct pathway neurons is gated by dopamine and excitatory synaptic input. Cell Rep. 43, (2024).

62. Ryan, M. B. Aberrant Striatal Activity in Parkinsonism and Levodopa-Induced Dyskinesia. Cell Rep. 23,**12**, 3438–3446 5 (2018).

63. Girasole, A. E. A Subpopulation of Striatal Neurons Mediates Levodopa-Induced Dyskinesia. Neuron 97,**4**, 787–795 6 (2018).

64. Fieblinger, T. Cell type-specific plasticity of striatal projection neurons in parkinsonism and L-DOPA-induced dyskinesia. Nat. Commun. 5 5316. 31, (2014).

65. Fieblinger, T. Striatal Control of Movement: A Role for New Neuronal (Sub-) Populations? vol. 15 697284 (2021).

66. Santamaria, F. The diffusional properties of dendrites depend on the density of dendritic spines. Eur. J. Neurosci. 34, 561–568 (2011).

67. Konradi, C. Transcriptome analysis in a rat model of L-DOPA-induced dyskinesia. Neurobiol. Dis. 17, 219–236 (2004).

68. Figge, D. A. Differential activation states of direct pathway striatal output neurons during L-DOPA-induced dyskinesia development. J. Neurosci. 44, e0050242024 (2024).

69. Chary, K. Diffusion MRI approaches for investigating microstructural complexity in a rat model of traumatic brain injury. Sci. Rep. 13, 2219 (2023).

70. Gittis, A. H., Hooks, B. M. & Gerfen, C. R. Basal ganglia circuits. in Neural Circuit and Cognitive Development 221–242 (Academic Press, 2020).

71. Fujita, M. Morphologic changes in striatonigral neurons in a rat model of levodopa-induced dyskinesia. Neurosci. Res. 104953 (2025).

72. Yoo, H. S. Cerebellar connectivity in Parkinson’s disease with levodopa-induced dyskinesia. Ann. Clin. Transl. Neurol. 6, 2251–2260 (2019).

73. Kishore, A. & Popa, T. Cerebellum in levodopa-induced dyskinesias: the unusual suspect in the motor network. Front. Neurol. 5, 157 (2014).

74. Coutant, B. Cerebellar stimulation prevents Levodopa-induced dyskinesia in mice and normalizes activity in a motor network. Nat. Commun. 13, 3211 (2022).

75. Koch, G. Cerebellar magnetic stimulation decreases levodopa-induced dyskinesias in Parkinson disease. Neurology 73, 113–119 (2009).

76. Hoshi, E. The cerebellum communicates with the basal ganglia. Nat. Neurosci. 8, 1491–1493 (2005).

77. Bostan, A. C., Dum, R. P. & Strick, P. L. The basal ganglia communicate with the cerebellum. in Proceedings of the national academy of sciences 107. 18 8452–8456 (2010).

78. Cardinale, A., De Iure, A. & Picconi, B. Neuroinflammation and dyskinesia: a possible causative relationship? Brain Sci. 14, 514 (2024).

79. Nimura, T. The role of σ-receptors in levodopa-induced dyskinesia in patients with advanced Parkinson disease: a positron emission tomography study. J. Neurosurg. 100, 606–610 (2004).

80. Goldman, J. G. Entorhinal cortex atrophy differentiates Parkinson’s disease patients with and without dementia. Mov. Disord. 27, 727–734 (2012).

81. Patel, R. Hippocampal and entorhinal cortex atrophy across the Parkinson’s disease cognitive impairment spectrum (S39. 004. Neurology 88, 39–004 (2017).

82. Camicioli, R. Parkinson’s disease is associated with hippocampal atrophy. Mov. Disord. Off. J. Mov. Disord. Soc. 18,**7**, 784–90 (2003).

83. Bouchard, T. P. Age and dementia-associated atrophy predominates in the hippocampal head and amygdala in Parkinson’s disease. Neurobiol. Aging 29,**7**, 1027–39 (2008).

84. Yang, J. Neuroprotective effect of curcumin on hippocampal injury in 6-OHDA-induced Parkinson’s disease rat. Pathol. Res. Pract. 210,**6**, 357–62 (2014).

85. Kim, J.-Y., Shim, J.-H. & Baek, H.-M. White matter microstructural alterations in newly diagnosed Parkinson’s disease: A whole-brain analysis using dMRI. Brain Sci. 12, 227 (2022).

86. Del Tredici, K. & Braak, H. Dysfunction of the locus coeruleus–norepinephrine system and related circuitry in Parkinson’s disease-related dementia. J. Neurol. Neurosurg. Psychiatry 84, 774–783 (2013).

87. Buhidma, Y. Potential of animal models for advancing the understanding and treatment of pain in Parkinson’s disease. NPJ Park. Dis. 6 **1**, (2020).

88. Domenici, R. A. Parkinson’s disease and pain: Modulation of nociceptive circuitry in a rat model of nigrostriatal lesion. Exp. Neurol. 315, 72–81 (2019).

89. Dauer, W. & Przedborski, S. Parkinson’s disease: mechanisms and models. Neuron 39, 889–909 (2003).

90. Cenci, M. A. & Lundblad, M. Ratings of L-DOPA-induced dyskinesia in the unilateral 6-OHDA lesion model of Parkinson’s disease in rats and mice. Curr. Protoc. Neurosci. 41, 9–25 (2007).

91. Allan Johnson, G. Whole mouse brain connectomics. J. Comp. Neurol. 527, 2146–2157 (2019).

92. Tendler, B. C. A method to remove the influence of fixative concentration on postmortem T2 maps using a kinetic tensor model. Hum. Brain Mapp. 42, 5956–5972 (2021).

93. Sun, S.-W. Relative indices of water diffusion anisotropy are equivalent in live and formalin-fixed mouse brains. Magn. Reson. Med. 50,**4**, 743–8 (2003).

94. Breese, G. R. & Traylor, T. D. Depletion of brain noradrenaline and dopamine by 6-hydroxydopamine. Br. J. Pharmacol. 42, 88–99 (1971).

95. Prasad, E. M. & Hung, S.-Y. Behavioral tests in neurotoxin-induced animal models of parkinson’s disease. Antioxidants 9, 1007 (2020).

96. Schallert, T. & Tillerson, J. L. Intervention Strategies for Degeneration of Dopamine Neurons in Parkinsonism: Optimizing Behavioral Assessment of Outcome. in Central Nervous System Diseases (eds. Emerich, D. F., Dean, R. L. & Sanberg, P. R.) 131–151 (Humana Press, Totowa, NJ, 2000). doi:10.1007/978-1-59259-691-1_8.

97. Yi, S. Y. Clinical translational neuroimaging of the antioxidant effect of N-acetylcysteine on neural microstructure. Magn. Reson. Med. 87,**2**, 820–836 (2022).

98. Johnson, G. A. A multicontrast MR atlas of the Wistar rat brain. NeuroImage 242, 118470 (2021).

99. Yeh, F.-C. Population-averaged atlas of the macroscale human structural connectome and its network topology. Neuroimage 178, 57–68 (2018).

100. Yeh, F.-C. Deterministic diffusion fiber tracking improved by quantitative anisotropy. PloS One 8, 80713 (2013).

101. Yeh, F.-C. Automatic Removal of False Connections in Diffusion MRI Tractography Using Topology-Informed Pruning (TIP. Neurotherapeutics 16, 52–58, (2019).

102. Yeh, F.-C. Quantifying differences and similarities in whole-brain white matter architecture using local connectome fingerprints. PLoS Comput. Biol. 12, (2016).

103. Rueden, C. T. ImageJ2: ImageJ for the next generation of scientific image data. BMC Bioinformatics 18, (2017).

