## Supplementary Figure and Table for "Levodopa induced dyskinesia severity is linked to distinct microstructural alterations in nigrostriatal and brainwide neural circuits"

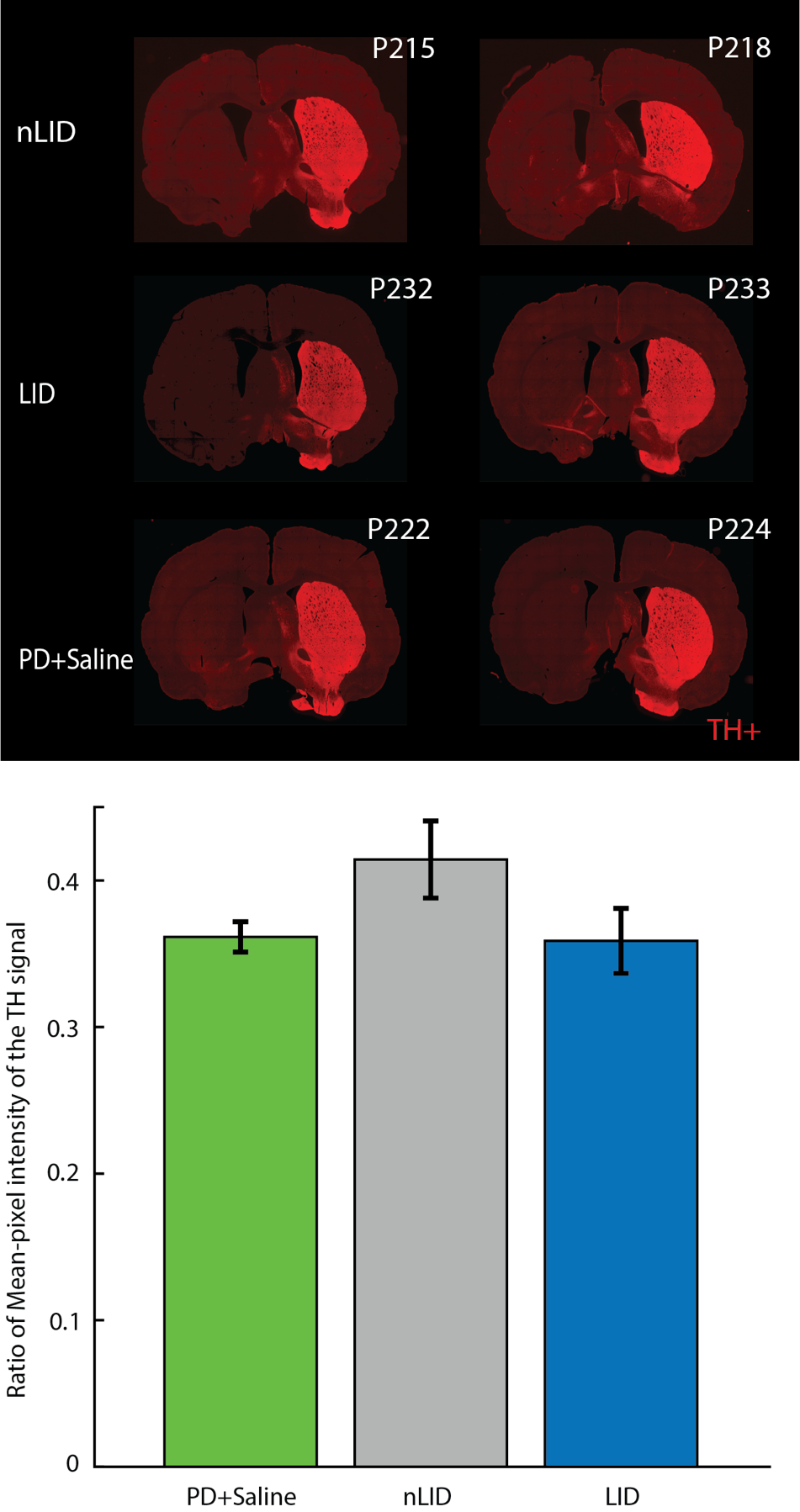

**Fig. S1 Representative histological sections from nLID, LID and PD+Saline.** Coronal striatal slices from 2 nLID, 2 LID and 2 PD+Saline; bright red regions mark dopaminergic neurons staining positive for tyrosine hydroxylase (TH+).

**Table 1: Histological and LID severity outcome for all rats completing the study.**

|  |  |  | L-R Striatal Intensity Ratio (AU) | Area Under AIMs Curve (AU) |
| --- | --- | --- | --- | --- |
| 1 | **LID** | P187 | 0.33 | 26 |
| 2 |  | P201 | 0.24 | 37 |
| 3 |  | P214 | 0.31 | 35 |
| 4 |  | P218 | 0.39 | 16 |
| 5 |  | p226 | 0.36 | 27 |
| 6 |  | p234 | 0.41 | 31 |
| 7 |  | p235 | 0.43 | 40 |
| 8 |  | p236 | 0.4 | 28 |
| 1 | **nLID** | p185 | 0.55 | 0 |
| 2 |  | p200 | 0.42 | 0 |
| 3 |  | p203 | 0.43 | 3 |
| 4 |  | p215 | 0.4 | 7 |
| 5 |  | p229 | 0.39 | 0 |
| 6 |  | p232 | 0.32 | 0 |
| 7 |  | p233 | 0.39 | 0 |
| 1 | **Saline** | p207 | 0.38 | 0 |
| 2 |  | p210 | 0.37 | 0 |
| 3 |  | p212 | 0.34 | 0 |
| 4 |  | p219 | 0.39 | 0 |
| 5 |  | p221 | 0.33 | 0 |
| 6 |  | p222 | 0.39 | 0 |
| 7 |  | p224 | 0.33 | 0 |
| 1 | **Vehicle** | p238 | 1.02 | 0 |
| 2 |  | p239 | 0.9 | 0 |
| 3 |  | p241 | 0.95 | 0 |
| 4 |  | p242 | 0.89 | 0 |
